# EG-BINN: Parameter-Efficient Ensembling of Gated Biologically Informed Neural Networks for Epigenetic Age Estimation

**DOI:** 10.64898/2026.10.17.759869

**Authors:** Travyse Edwards, Qi Long, Li Shen

## Abstract

Epigenetic aging clocks are widely used to evaluate biological aging, disease risk, and clinical interventions. However, most existing clocks rely on linear models that overlook nonlinear relationships between age and methylation patterns, or on deep learning architectures that are parameter-intensive and difficult to interpret. We developed EG-BINN, a parameter-efficient ensemble of biologically informed neural networks that uses sample-specific gating to modulate representations across biologically annotated nodes. Trained on 93 multi-tissue datasets, EG-BINN achieved the lowest mean and median absolute errors among matched linear and nonlinear baselines in five-fold cross-validation, as well as the lowest mean absolute and root mean squared errors across eight held-out Gene Expression Omnibus (GEO) datasets. Analysis of the gating mechanism revealed widespread associations between pathway gate values and age, including trajectories that varied across blood, adipose tissue, skeletal muscle, and liver. Across 11 TCGA cohorts, EG-BINN identified significant tumor–normal differences in residual epigenetic age acceleration in six cohorts, whereas Shapley additive explanations (SHAP) highlighted recurrent pathway nodes contributing to tumor age predictions. EG-BINN combines accurate pan-tissue age prediction with an explainable architecture for examining biologically labeled nodes across age, tissue, and disease contexts.

## Introduction

Nearly 22% of the global population is projected to be 60 years of age or older by 2050 [1]. This demographic shift has major implications for healthcare, as chronological age is one of the strongest risk factors for a wide range of chronic diseases, including cancer, cardiovascular disease, and neurodegenerative disorders. However, differences in health status and disease risk among individuals of the same age suggest that chronological age alone does not adequately reflect the heterogeneity of the aging process. This limitation has motivated the development of biological age as a quantitative measure intended to better reflect an individual’s physiological condition and disease risk. In recent years, numerous models designed to estimate biological age, commonly referred to as aging clocks, have been developed using a wide range of molecular and physiological biomarkers. Among these, DNA methylation (DNAm)-based biomarkers have emerged as some of the most robust indicators of biological aging, making epigenetic clocks one of the most extensively studied classes of aging biomarkers.

Traditional epigenetic aging clocks, often referred to as first-generation clocks, typically train penalized regression models on DNAm measurements from healthy individuals to model normative aging. When applied to populations of mixed or unknown health, deviations between predicted and true chronological age are interpreted as accelerated or decelerated biological aging. The 353-CpG Horvath pan-tissue clock is an example of a highly influential first-generation clock, which demonstrated accurate age prediction across a broad range of human tissues and cell types [2]. Subsequent efforts led to the development of second-generation clocks, which were trained to predict mortality risk, healthspan-related phenotypes, or composite measures of physiological decline, thereby capturing biological aging more directly than chronological age alone [3–6].

Despite their success, many of these previous models have several limitations. First, accumulating evidence suggests that many age-associated methylation changes exhibit nonlinear trajectories across the lifespan, which may not be fully captured by the aforementioned linear models [7, 8]. Second, although these models often have the benefit of interpretability, their utility for broader biological discovery may be constrained by their reliance on relatively small subsets of CpG sites. Many linear epigenetic clocks incorporate only a few hundred CpGs, despite modern DNA methylation platforms profiling hundreds of thousands of sites across the genome. Consequently, much of the measured methylome does not contribute directly to the final predictive model. While this sparsity can support robust age prediction, it may also restrict the identification of distributed aging-associated methylation patterns and the broader gene- and pathway-level processes underlying epigenetic aging. These limitations motivate approaches that leverage a substantially larger fraction of the methylome while capturing complex patterns of age-associated methylation.

Several deep learning approaches for epigenetic age prediction have been developed to address these gaps [9–12]. Among these, AltumAge demonstrated that neural networks, such as the densely connected multilayer perceptron (MLP), can effectively model age using more than 20,000 CpG sites, substantially expanding feature coverage beyond that of prior penalized regression-based clocks [11]. However, scaling these densely connected architectures to accommodate the hundreds of thousands of CpGs measured by modern arrays substantially increases the number of trainable parameters and the risk of overfitting, especially given the relatively limited sample sizes typically used for aging clock development.

One promising strategy for addressing this challenge is the use of biologically informed neural networks (BINNs), which incorporate prior knowledge of gene and pathway relationships directly into the network architecture. By imposing biologically motivated connectivity constraints, BINNs reduce the number of trainable parameters through structured sparsity while improving model interpretability [13]. However, existing BINNs generally use the same biologically defined relationships to aggregate information across samples, even though the relative contributions of molecular signals to gene and pathway representations may differ across tissues and molecular contexts [14–16]. Conventional BINNs may capture such variation through input-dependent activations, but they lack an explicit inductive bias for conditionally modulating biologically annotated representations. Multiplicative interactions, including gating, have been proposed as a useful inductive bias for conditional computation and the integration of contextual information [17]. BINNs may also exhibit variability across training runs, potentially affecting the stability of their predictions. Although independently trained ensembles can mitigate this variability, they increase the computational cost and complexity of model training.

To address these limitations, we developed EG-BINN, a framework that combines parameter-efficient ensembling with gated, biologically informed neural networks for pan-tissue age prediction. EG-BINN was trained on 189,291 CpG sites measured using the Illumina HumanMethylation450K and MethylationEPIC platforms (Fig. 1). Gating allows gene and pathway nodes to be modulated according to sample context [18, 19], while parameter-efficient ensembling captures multiple learned representations without requiring several independently parameterized networks [20]. Both components operate within the structured biological hierarchy of the BINN architecture. EG-BINN achieved strong predictive performance and cross-tissue generalization relative to pre-existing epigenetic clocks and neural network baselines. Analysis of the learned gating patterns identified pathway representations whose modulation varied with age and tissue context. SHAP analysis of these pathway representations further revealed recurrent contributors to age predictions, as well as representations that were differentially prioritized in tumor samples relative to cohort-specific normal reference distributions. Together, these findings show that biologically informed deep learning can support accurate age prediction while providing structured, hypothesis-generating insights into molecular representations associated with age, tissue context, and disease.

**Figure 1:**
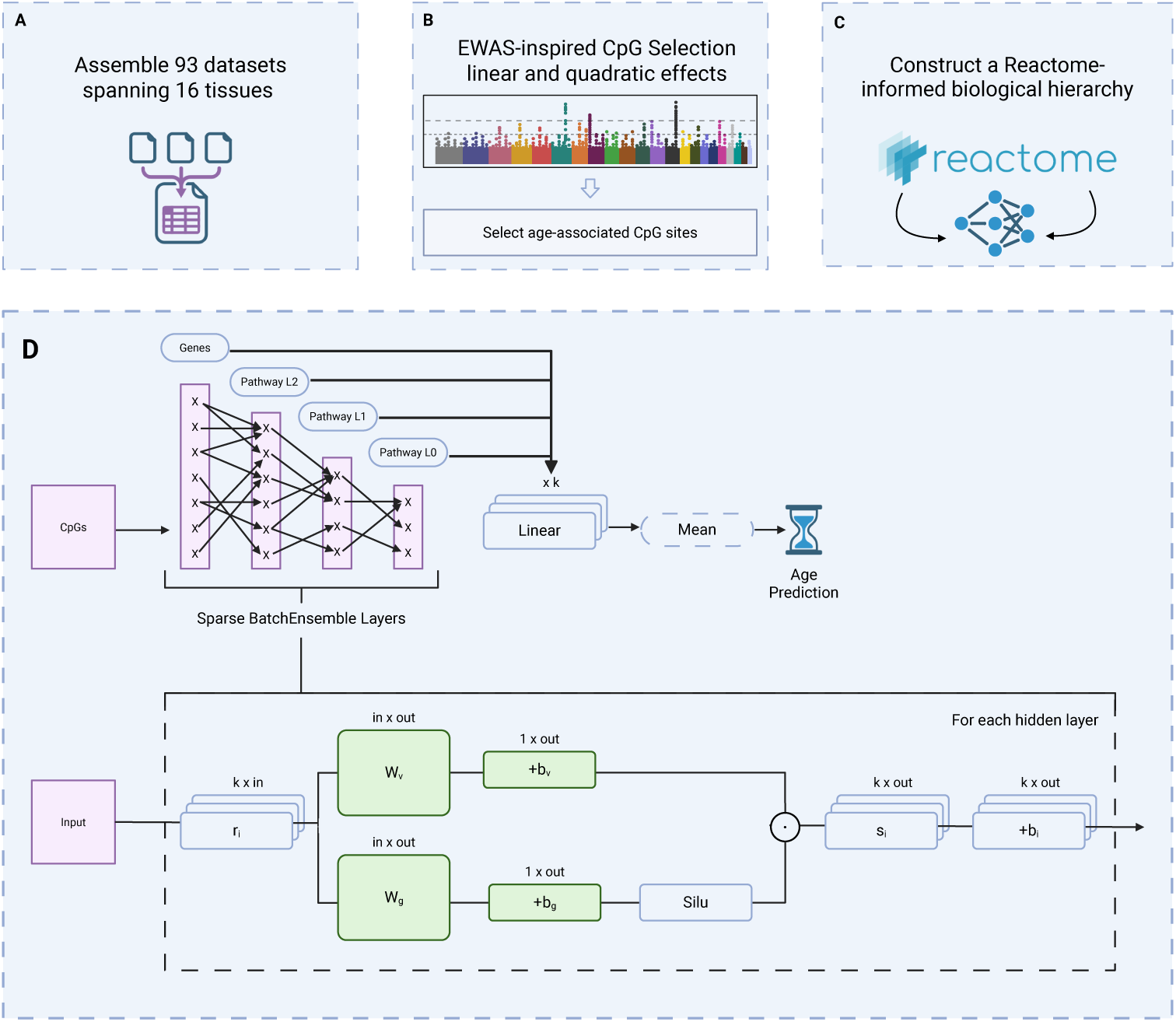
Development and architecture of EG-BINN. **a**, DNA methylation data were assembled from 93 Gene Expression Omnibus datasets spanning a variety of tissues and cell types. **b**, Age-associated CpG sites were selected using linear and quadratic age terms, with feature selection restricted to training data. **c**, Selected CpGs were mapped to genes and organized into a hierarchical biological structure using Reactome pathway annotations. **d**, EG-BINN connects CpGs, genes, and successive pathway levels through biologically constrained sparse layers. Each hidden layer incorporates sample-specific gating, and parameter-efficient ensemble members share core model parameters while retaining member-specific adaptations. Prediction heads at multiple hierarchy levels provide multistage supervision, and their outputs are aggregated across hierarchy levels and ensemble members to produce the final age estimate.

## Results

### Model evaluation and architectural ablation

BINNs assign hidden nodes to specific genes or pathways and restrict their connections according to known biological relationships, creating a sparse alternative to a densely connected neural network. These models generally apply the same learned transformations across samples, even though the contributions of individual genes and pathways to age prediction may vary with age, tissue type, and other biological factors. They also inherit the sensitivity of neural networks to random initialization, which can lead to variation in performance across trained models. EG-BINN addresses these limitations in two ways. First, sample-specific gating controls how much information from hidden nodes is passed through the biological hierarchy. Second, TabM-based ensembling combines shared layer weights with lightweight member-specific adaptations, allowing distinct age predictions to be generated and averaged across ensemble members without requiring several independently parameterized networks (Fig. 1). Together, these components allow biologically organized information to be modulated across samples while reducing reliance on any single model initialization.

We first compared EG-BINN with matched linear, dense MLP, and traditional BINN baselines. Each model was optimized and evaluated using the same five dataset-level cross-validation folds. The configuration with the best cross-validation performance for each model is reported in Table 1.

**Table 1:**
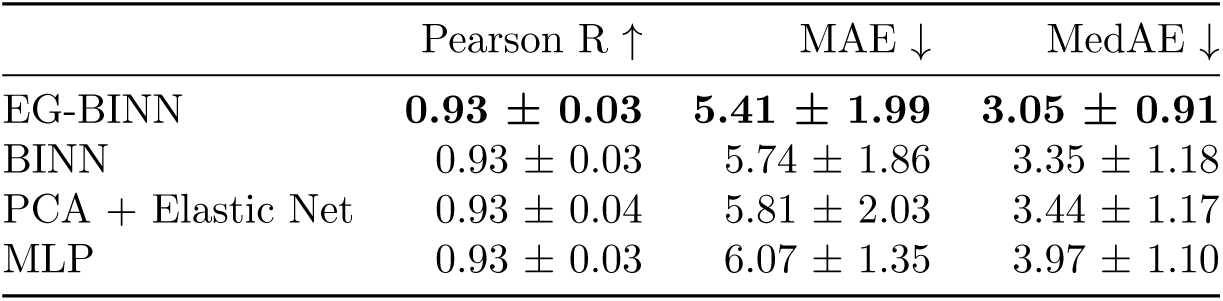
Predictive performance across five-fold cross-validation

EG-BINN achieved the lowest prediction error, with a mean absolute error (MAE) of 5.41 *±* 1.99 years and a median absolute error (MedAE) of 3.05 *±* 0.91 years. BINN and PCA + Elastic Net achieved MAEs of 5.74 *±* 1.86 and 5.81 *±* 2.03 years, respectively, while the MLP had the highest MAE at 6.07 *±* 1.35 years. Pearson correlations were similar across models, with mean values of approximately 0.93. The best-performing hyperparameter configuration identified for each model was then used to retrain that model on the complete training data. These final models, together with previously published clocks, were evaluated on eight held-out GEO datasets that had not been used for feature selection, model selection, or hyperparameter optimization. EG-BINN achieved an MAE of 3.39 years, a MedAE of 2.22 years, an RMSE of 5.28 years, and a Pearson correlation of 0.88 (Table 2). Among all evaluated models, EG-BINN had the lowest MAE and RMSE and tied with ZhangEN for the highest Pearson correlation. ZhangBLUP achieved a slightly lower MedAE than EG-BINN (2.12 versus 2.22 years), although its MAE and RMSE were higher at 3.52 and 6.19 years, respectively. Overall, EG-BINN combined the lowest average prediction error and RMSE with one of the strongest correlations between predicted and chronological age across the held-out datasets.

**Table 2:** Predictive performance on the eight-dataset held-out benchmark

| Model | Required CpG inputs | Pearson R ↑ | MAE ↓ | MedAE ↓ | RMSE ↓ |
| --- | --- | --- | --- | --- | --- |
| EG-BINN | 189,291 | **0.88** | **3.39** | <u>2.22</u> | **5.28** |
| PCA + Elastic Net | 189,291 | <u>0.84</u> | 4.07 | 2.79 | <u>6.09</u> |
| ZhangBLUP | 319,607 | <u>0.84</u> | <u>3.52</u> | **2.12** | 6.19 |
| BINN | 189,291 | 0.83 | 4.19 | 2.79 | 6.34 |
| ZhangEN | 514 | **0.88** | 5.08 | 3.68 | 7.42 |
| MLP | 189,291 | 0.78 | 4.85 | 3.08 | 7.55 |
| PCHannum | 78,464 | 0.77 | 8.34 | 7.58 | 10.22 |
| Hannum | 71 | 0.59 | 8.83 | 7.99 | 10.76 |
| YingAdaptAge | 999 | 0.52 | 14.92 | 7.84 | 21.61 |
| YingDamAge | 1,089 | 0.06 | 16.44 | 6.32 | 24.72 |
| YingCausAge | 585 | 0.02 | 27.48 | 10.48 | 39.08 |

We next evaluated how ensemble size, layer type, and supervision strategy affected predictive performance while holding all other model and training settings fixed to those of the final EG-BINN configuration (Supplementary Table S1). The configuration combining eight ensemble members, gated layers, and multistage supervision performed best across all reported metrics, achieving a

Pearson correlation of 0.86 *±* 0.01, an MAE of 4.03 *±* 0.10 years, a MedAE of 2.51 *±* 0.10 years, and an mean squared error (MSE) of 38.61 *±* 1.96 years^2^. Each component improved performance when the other two were held fixed. Reducing the ensemble from eight members to one increased the MAE from 4.03 to 4.10 years. Removing the gating from each hidden layer increased the MAE to 4.26 years and reduced the Pearson correlation from 0.86 to 0.83. Restricting supervision to the final output produced the largest individual loss in performance, increasing the MAE to 4.45 years and reducing the Pearson correlation to 0.83. Performance declined further when final-output supervision was combined with sparse layers or a single model, with the single-model sparse configuration producing the highest MAE of 5.39 *±* 0.34 years.

### Age- and tissue-dependent pathway gating

To examine how gating varied with age and whether these relationships differed among tissues, we analyzed gate values in the first pathway-level layer (L2; Fig. 1d). Linear regression models adjusted for tissue and study identified significant age associations for 423 of the 504 L2 pathway nodes after Benjamini–Hochberg correction (Fig. 2a). The strongest positive associations included Signaling by Rho GTPases, Neurexins and neuroligins, and Regulation of lipid metabolism by PPAR*α*, whereas Formation of the ureteric bud, Generic Transcription Pathway, and other developmental and transcription-related pathways showed some of the strongest negative associations. Most displayed gate values remained above the neutral value of one across the observed age range; positive associations therefore reflected increasing amplification of the corresponding pathway-level output, whereas most negative associations reflected decreasing amplification rather than attenuation. Ageby-tissue interaction models adjusted for study further identified significant differences among blood, adipose tissue, skeletal muscle, and liver (Fig. 2b). Among pathways with significant interactions after Benjamini–Hochberg correction, Signaling by Rho GTPases, Fatty acid metabolism, Neurexins and neuroligins, Viral Infection Pathways, and Signaling by interleukins showed the largest differences in tissue-specific age slopes, involving the magnitude and, for some pathways, the direction of the association.

**Figure 2:**
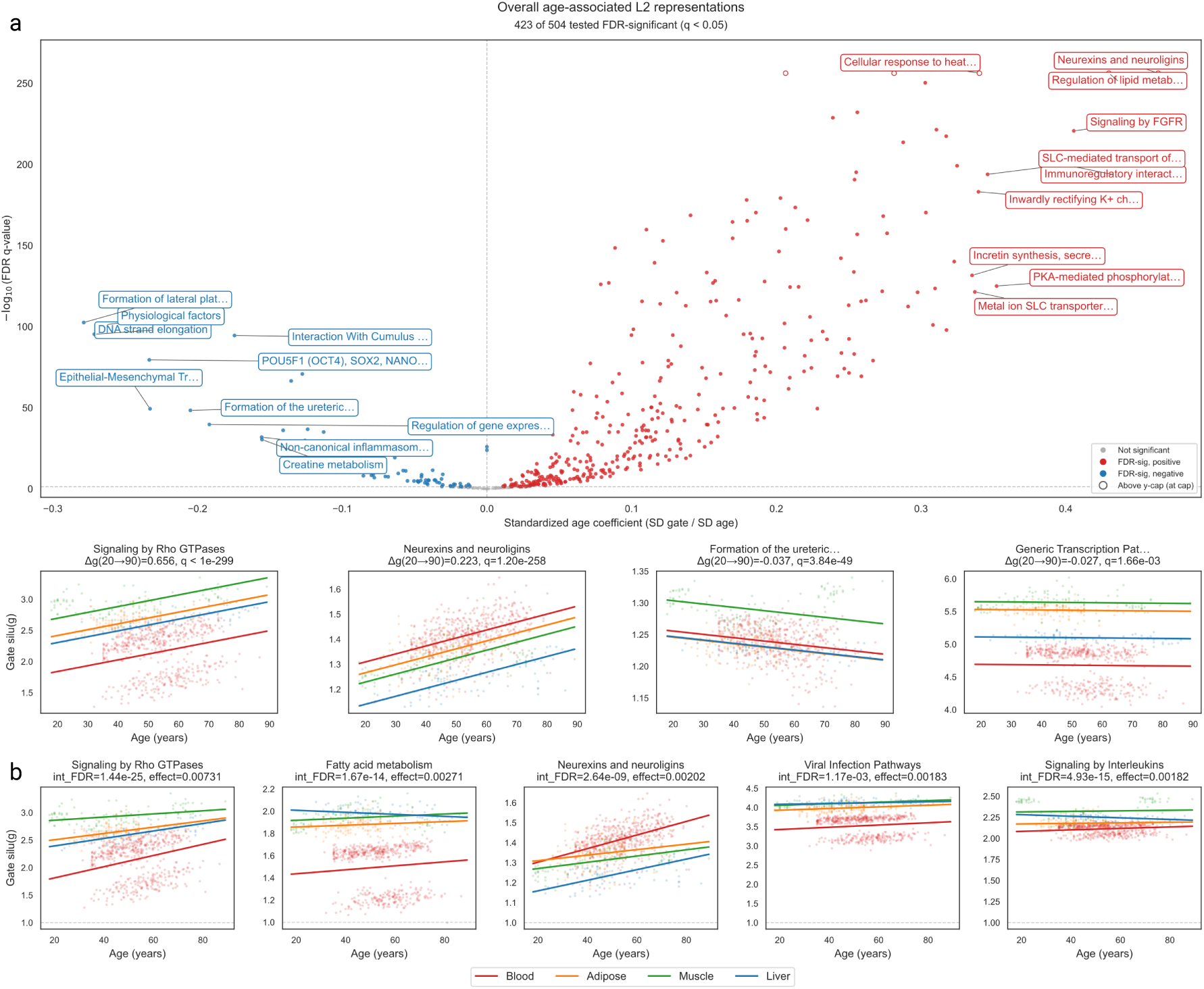
Age- and tissue-dependent modulation of pathway representations in EG-BINN. **a**, Standardized associations between chronological age and gating values for 504 L2 pathway representations, estimated across blood, adipose tissue, skeletal muscle, and liver with adjustment for tissue and study. Filled points indicate associations significant after Benjamini–Hochberg correction. Trajectory plots show adjusted gating values for the significant representations with the largest positive and negative estimated changes from ages 20 to 90. **b**, Adjusted tissue-specific trajectories for five L2 representations with significant age-by-tissue interactions and the largest differences among tissue-specific age slopes. Lines show fitted values, and shaded bands indicate 95% confidence intervals. The horizontal dashed line at a gate value of 1 indicates neutral modulation; values above 1 indicate amplification, whereas values between 0 and 1 indicate attenuation of the corresponding representation.

### Biological Application of EG-BINN

Given the association between epigenetic age acceleration and age-related disease observed in literature, we applied EG-BINN to tumor and normal samples from 11 cohorts in The Cancer Genome Atlas (TCGA). We first evaluated age-prediction performance in normal tissues. We then compared residual epigenetic age acceleration between tumor and normal samples and identified the pathway nodes contributing to shifts in age predictions for tumor samples relative to cohort-specific normal backgrounds. AltumAge and Horvath’s pan-tissue clock were excluded from these comparisons because TCGA samples were included in their training data.

#### Age prediction and residual age acceleration in TCGA

We first evaluated model performance in normal tissues from 11 TCGA cohorts (Fig. 3a). EG-BINN achieved the lowest average cohort-level MedAE among the evaluated models, with a mean of 6.8 *±* 2.9 years across cohorts. BINN and PCA + Elastic Net had the next-lowest average MedAEs at 7.8 *±* 3.5 and 8.0 *±* 3.8 years, respectively. Next, we compared residual epigenetic age acceleration between tumor and normal samples within each cohort (Fig. 3b). After Benjamini–Hochberg correction across the 11 cohort-level comparisons, six cohorts showed significant tumor–normal differences. Residual epigenetic age acceleration was higher in tumor samples from LIHC, KIRC, THCA, LUAD, and BRCA, but lower in UCEC tumors. No significant tumor–normal differences were observed in COAD, PRAD, KIRP, HNSC, or LUSC.

**Figure 3:**
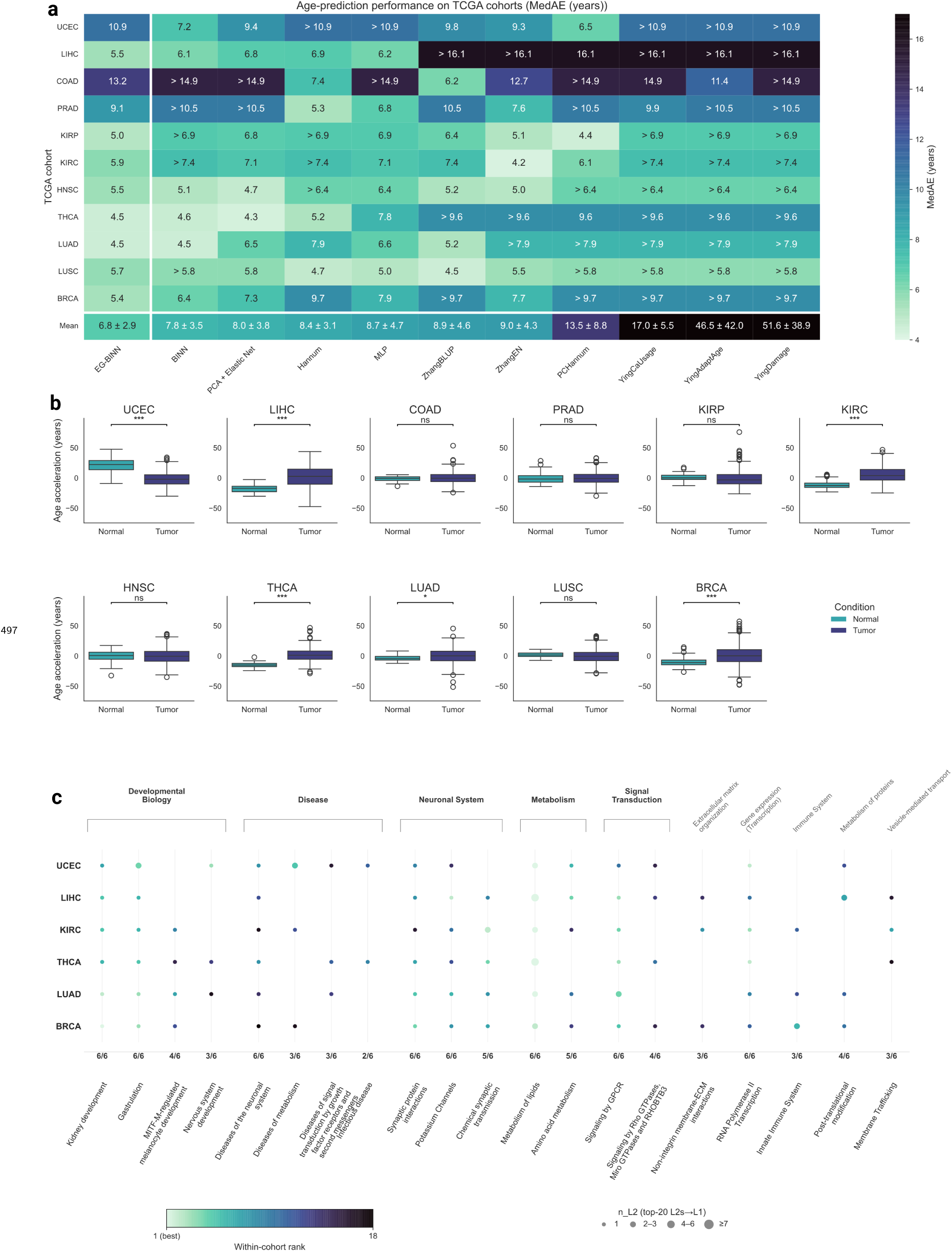
Evaluation of EG-BINN in TCGA normal and tumor tissues. **a**, MedAE of EG-BINN and comparator models in normal samples from 11 TCGA cohorts. Columns are ordered by increasing mean MedAE across cohorts, and the summary row shows the unweighted mean *±* standard deviation of the cohort-level values. **b**, Residual epigenetic age acceleration in normal and tumor samples within each cohort. Groups were compared using two-sided Mann–Whitney *U* tests with Benjamini–Hochberg correction across cohorts. Boxes show the median and interquartile range, and whiskers extend to 1.5 times the interquartile range. **c**, Recurrent L2 pathway representations among the 20 highest-ranked representations by mean absolute SHAP value in each of the six cohorts with significant tumor–normal differences in residual epigenetic age acceleration. Columns show representations retained in at least three cohorts. Dot color indicates within-cohort rank, and values below each column indicate the number of cohorts in which the representation was retained. Missing dots indicate representations outside the corresponding cohort’s top 20.

#### Pathway attribution across TCGA cohorts

We next used SHAP to identify the L2 pathway nodes that contributed most strongly to age predictions for tumor samples, using normal samples from the same cohort as the background reference (Fig. 3c). Ranking nodes within each cohort by mean absolute attribution revealed a recurrent set of 12 pathway representations ranked among the top 20 in at least five of the six cohorts, alongside nodes with cohort-variable rankings. Eight nodes ranked in the top 20 across all six cohorts, encompassing receptor and retinoic acid signaling (Downstream signaling of activated FGFR, GPCR downstream signaling, and Signaling by Retinoic Acid), synaptic organization (Neurexins and neuroligins and Long-term potentiation), lipid metabolism (Fatty acid metabolism), basal transcription (Generic Transcription Pathway), and chaperonin folding (Formation of tubulin folding intermediates by CCT/TriC). Four additional metabolic, signaling, and matrix-related nodes ranked in the top 20 in five cohorts (Metabolism of vitamins and cofactors, RHO GTPases Activate PKNs, Keratan sulfate biosynthesis, and Neurotransmitter receptors and postsynaptic signal transmission). Notably, three universally recurrent nodes (Fatty acid metabolism, Neurexins and neuroligins, and Generic Transcription Pathway) were also highlighted in the held-out dataset gating analysis (Fig. 2), linking the L2 representations identified by both analyses.

## Discussion

In this study, we developed EG-BINN, a biologically informed architecture for pan-tissue epigenetic age prediction. EG-BINN improved prediction relative to matched linear and nonlinear baselines in cross-validation and achieved the lowest MAE and RMSE among the models evaluated on the held-out benchmark. Ablation analyses supported the combined use of gated layers, multistage supervision, and parameter-efficient ensembling. Analysis of the gating mechanism in the held-out datasets revealed widespread associations between chronological age and the gate values of L2 pathway representations, with a subset exhibiting tissue-dependent trajectories. Evaluating EG-BINN in TCGA showed favorable, but cohort-dependent predictive performance. Residual epigenetic age acceleration differed significantly between tumor and normal samples in six of eleven cohorts, while SHAP analysis identified recurrent and cohort-dependent L2 pathway representations contributing to tumor-sample predictions. Together, these findings show that EG-BINN combines strong pan-tissue predictive performance with biologically annotated internal representations that can be examined across age, tissue, and disease contexts.

One of the core benefits of EG-BINN lies at the intersection of intrinsic interpretability and post-hoc explainability. Although the definition of these terms can vary in the literature, interpretability is commonly used to describe models whose internal operation is understandable from their structure, whereas explainability often refers to methods applied after training to characterize what influences a model’s predictions [21, 22]. EG-BINN retains the biologically annotated nodes and constrained connectivity that provide intrinsic interpretability in conventional BINNs, while sample-specific gating provides an explicit inductive bias allowing the gene and pathway representations to be modulated before contributing to age predictions. The widespread gate–age associations indicate that gating was used dynamically rather than functioning as an approximately static scaling operation. The tissue interactions further show that the shared hierarchy was not used identically across tissues, supporting the intended role of gating as a mechanism for context-dependent modulation.

Biologically informed architectures also provide a means of explaining epigenetic age predictions at the gene and pathway levels. Previous deep learning-based epigenetic clocks, including AltumAge [11], DeepMAge [9], MethylNet [10], and DeepStrataAge [12], have used post-hoc methods to identify CpGs that contribute to age predictions. Interpreting these findings at the pathway level often requires mapping highly attributed CpGs to genes and performing enrichment analysis. However, correlations among CpGs can complicate the assignment of individual feature importance [21], while selecting only highly ranked CpGs may overlook pathways whose predictive contributions are distributed across many features. By incorporating biological relationships directly into the network architecture, BINNs allow predictive contributions to be evaluated at the gene and pathway levels rather than inferred solely from individual CpG rankings [13, 23, 24]. This provides a more direct link between the model’s predictions and established biological pathways, although interpretation remains dependent on the underlying annotations. This framework allowed tumor-sample predictions in TCGA to be attributed directly to biologically annotated pathway representations. Highly attributed L2-level pathways recurrent across cancer types belonged to broad functional domains related to development, metabolism, neuronal signaling, disease, and signal transduction. Their recurrence is consistent with shared features of cancer biology, including altered differentiation, epigenetic regulation, cellular signaling, and metabolism [25–27].

Several limitations should be acknowledged. First, the training data were dominated by blood and, to a lesser extent, brain, with fewer samples representing other tissues. This imbalance may affect generalization and calibration in tissues that were underrepresented during training. Second, we did not explicitly harmonize batch effects across the training and inference datasets. We incorporated a large number of independent datasets to reduce reliance on the characteristics of any single study. However, differences in sample processing, preprocessing, and study composition may remain, and their relationships with tissue representation and demographic composition could affect model performance in ways that cannot be separated in the current analysis. Third, the model architecture depends on the annotations used to map CpGs to genes and organize genes within the Reactome hierarchy [28]. The resulting representations therefore inherit the coverage and curation biases of these resources. Moreover, age-predictive methylation signals are not limited to CpGs with gene annotations, and excluding unmapped CpGs may remove information relevant to age prediction. Fourth, although our analyses establish that EG-BINN gating varies systematically with age and tissue, we did not directly examine tissue-dependent internal representations within the conventional BINN. We therefore cannot determine the extent to which gating captures tissue context beyond the tissue-dependent variation that can already arise as different inputs propagate through the conventional architecture. This is a worthwhile direction to explore in the future to better understand the architecture. Finally, inconsistent metadata prevented systematic evaluation of model performance across sex, ancestry, race and ethnicity, and other population characteristics, limiting our ability to assess generalizability across demographic groups.

Future work could extend EG-BINN to larger DNA methylation assays, such as the Human-MethylationEPIC array, to increase CpG, gene, and pathway coverage. Training on more balanced tissue collections and diverse populations would also enable a more comprehensive evaluation of model performance and generalizability. Evaluating associations with age-related disease, functional decline, mortality, and intervention response will be important for determining whether residual EG-BINN age captures biologically relevant variation beyond chronological age prediction. The architecture could also be adapted to transcriptomic or proteomic data, either to develop aging predictors within individual molecular modalities or to integrate multiple data types within a multiomics framework. Overall, EG-BINN demonstrates that biologically informed neural networks can support accurate pan-tissue epigenetic age prediction while providing a structured view of how age-predictive methylation information is modulated across age, tissue, and disease contexts.

## Methods

### Cohorts and Data Sources

DNAm data were obtained from publicly available studies in GEO and TCGA [29, 30]. The GEO collection comprised 18,329 samples from 101 datasets. Of these, 13,898 samples from 93 datasets formed the training collection, and 4,431 samples from eight additional datasets were reserved for held-out evaluation. TCGA samples from the UCEC, LIHC, COAD, PRAD, KIRP, KIRC, HNSC, THCA, LUAD, LUSC, and BRCA cohorts were used separately for downstream analyses of normal and tumor tissues. Dataset characteristics for the 93 training datasets and eight held-out datasets are provided in Supplementary Fig. S1 and Supplementary Dataset S1. Dataset information for TCGA can be found in Supplementary Dataset S2.

#### DNAm Data Processing

All analyses used publicly available, source-processed DNAm beta-value matrices. Upstream processing varied across studies and is documented in the corresponding GEO records and publications. No additional global normalization or cross-study batch correction was performed. Beta values were used as model inputs, while M-values were used exclusively for PCA- and epigenome wide association study (EWAS)-based feature selection.

Quality control was performed independently for each GEO dataset. Samples and CpGs with more than 20% missing values were excluded. Following Horvath’s quality-control approach [2], samples were additionally excluded if their mean inter-array correlation, defined as the average correlation with the remaining samples in the dataset, was *≤* 0.99 or their maximum beta value across all probes was *≤* 0.96. After sample filtering, CpG missingness was recalculated, and sites exceeding the 20% threshold were removed. Only autosomal probes with identifiers beginning with cg were retained.

Missing beta values were imputed using CpG-wise training-set means. CpGs present in the training feature set but absent from validation or external datasets (held-out GEO or TCGA) were introduced during feature alignment and assigned their corresponding training-set means. For neural-network models, beta values were standardized using training-set CpG-wise means and standard deviations, then clipped to [*−*5, 5].

To prevent data leakage, all data-dependent preprocessing, including imputation, standardization, feature selection, and PCA fitting, was performed exclusively on the training partition within each cross-validation fold. The resulting transformations were applied to validation data without refitting. Following model and hyperparameter selection, preprocessing parameters were re-estimated on the complete development collection and applied unchanged to the held-out GEO and TCGA datasets.

### CpG feature selection and gene annotation

We performed an EWAS to identify CpG sites with linear or quadratic associations with chronological age and reduce the dimensionality of the methylation data before model training. Feature selection began with 453,054 CpGs that passed quality control. Within each training partition, missing beta values were imputed using CpG-wise means calculated from the training data, and the resulting values were logit-transformed to M-values.

For each CpG, separate ordinary least squares regression models were fitted with mean-centered chronological age as the dependent variable and either the CpG M-value alone or both its linear and quadratic terms as independent variables (Age *∼* CpG and Age *∼* CpG +CpG^2^, respectively). Both models were adjusted for the first 20 principal components (PCs) of the methylation M-value matrix, following Bernabeu et al. [3]. PCs were calculated using incremental principal component analysis (IncrementalPCA), fitted independently within each training partition using 500 components and a batch size of 1,024. M-values were centered but not scaled before PCA. Associations were assessed using two-sided *t*-tests of the linear CpG coefficient in the first model and the quadratic coefficient in the second model. CpGs meeting a Bonferroni-corrected significance threshold of *P <* 0.05*/N*_CpG_, where *N*_CpG_ denotes the number of tested CpGs, were retained. The final feature set comprised the union of CpGs identified by either model.

Feature selection was performed independently within each cross-validation training partition to prevent information leakage into validation data. Following model and hyperparameter selection, the procedure was repeated using the complete development collection to obtain the CpG set for final model training and held-out evaluation. Selected CpGs were then mapped to genes using the geneSymbol annotations in the Illumina Infinium MethylationEPIC manifest. CpGs without an annotated gene were excluded, while CpGs annotated to multiple genes were connected to each corresponding gene. These associations defined the CpG-to-gene connections in the first layer of the biologically informed network.

### Model architecture

EG-BINN was constructed as a biologically informed neural network organized according to the Reactome pathway hierarchy. Information propagated from CpG sites to genes and then through three successive pathway levels:

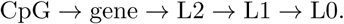

CpG-to-gene connections were derived from the Illumina Infinium MethylationEPIC manifest, as described above. Gene-to-pathway assignments and relationships among pathway levels were obtained from the Reactome Pathway Database. Reactome annotations and hierarchy relationships were downloaded from the current release endpoint on June 26, 2026, restricted to *Homo sapiens*, and mapped using Ensembl gene identifiers. L2 represented the first pathway layer above genes, while L1 and L0 contained successively broader pathway groupings.

Feature selection using the complete training data identified 267,791 CpGs associated with chronological age. Of these, 189,291 mapped to at least one gene in the biological hierarchy and were used as inputs to EG-BINN. The same CpG feature set was supplied to the BINN and MLP to support matched comparisons. Because a CpG could map to more than one gene, the selected features formed 209,839 CpG-to-gene edges. The resulting full-data network contained 189,291 CpG inputs, 20,031 gene nodes, 504 L2 pathway nodes, 132 L1 pathway nodes, and 28 L0 root pathway nodes. The L0 nodes represented broad biological domains, including Metabolism, Immune System, and Hemostasis.

Each hidden layer contains a value branch and a gating branch. Given an input representation **x**, these branches were defined as

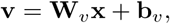

and

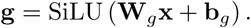

The output of the layer was then calculated as

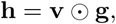

where *⊙* denotes elementwise multiplication. Both the value and gating weight matrices followed the sparse connectivity pattern defined for the corresponding biological layer. A gate value of 1 left the corresponding value-branch output unchanged, values greater than 1 increased its magnitude, values between 0 and 1 reduced its magnitude, and negative values reversed its sign. Gate biases were initialized to the value at which the SiLU activation equals one (*≈* 1.28), with gate weights scaled towards zero, so that each gated layer behaved as an approximately identity modulation at the start of training and learned to deviate as a function of sample context.

EG-BINN additionally incorporated a parameter-efficient ensemble inspired by TabM, with *K* = 8 members. Rather than training eight independently parameterized networks, the ensemble shared the biologically constrained value- and gating-branch weights across members. Ensemble diversity was introduced through member-specific input-scaling vectors, output-scaling vectors, additive biases, and prediction-head parameters. Each member therefore learned a distinct adaptation of the shared biological network without duplicating its sparse base weight matrices.

To provide supervision throughout the biological hierarchy, a prediction head was attached to the output of each hidden layer: gene, L2, L1, and L0. The model therefore contained *H* = 4 hierarchy-level heads, each of which generated a separate age prediction for every ensemble member. For a batch of *B* samples, the resulting prediction tensor had shape (*B, H, K,* 1). During training, the Huber loss with *δ* = 0.36 was calculated for every head and ensemble member and averaged equally across samples, members, and hierarchy levels:

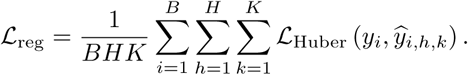

All four heads remained active during inference. The final prediction for sample *i* was the unweighted mean across hierarchy levels and ensemble members:

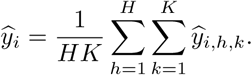

### Model Selection and Hyperparameter optimization

EG-BINN was compared with PCA + Elastic Net, an MLP, and a standard BINN using the same five dataset-level cross-validation folds. Model performance was evaluated using mean absolute error (MAE), median absolute error (MedAE), and Pearson correlation. Root mean squared error (RMSE) was additionally reported for the held-out benchmark.

Hyperparameters were optimized separately for each model using Optuna’s Tree-structured Parzen Estimator (TPE) sampler for 100 trials. Each trial evaluated one hyperparameter configuration across all five cross-validation folds. Models were trained for up to 101 epochs per fold using early stopping with a patience of 30 epochs for the neural network based models. All neural network-based models used the AdamW optimizer, a cosine-annealing learning-rate schedule without warmup, and a batch size of 256.

For each fold, model performance was measured by pooling all validation samples across the datasets assigned to that fold and calculating a single MSE. The five fold-specific MSE values were then averaged to obtain the optimization objective for the trial. To ensure a matched comparison within each fold, all three models received the same inputs selected from that fold’s training partition. After model and hyperparameter selection, feature selection was repeated using the complete development collection, yielding the 189,291 CpG measurements used for final training and held-out evaluation.

PCA + Elastic Net served as the linear baseline. Within each cross-validation fold, the standardized, gene-mapped CpGs selected from that fold’s training partition were reduced to 1,000 principal components using IncrementalPCA with a batch size of 1,000. The number of components was fixed rather than optimized. PCA was fitted independently within each cross-validation training fold and applied without refitting to the corresponding validation fold. Optuna was used to optimize the regularization strength *α*, while the mixing parameter was fixed at an *ℓ*_1_ ratio of 0.5.

For EG-BINN, the learning rate (2 *×* 10*^−^*^3^ to 6 *×* 10*^−^*^3^; log-uniform) and Huber loss delta (0.025 to 1.0; log-uniform) were optimized, while the ensemble size was fixed at *K* = 8. For the MLP baseline, the learning rate (2 *×* 10*^−^*^3^ to 6 *×* 10*^−^*^3^; log-uniform), hidden-layer width (*{*16, 32, 64, 128*}*), number of hidden layers (*{*2, 4, 6*}*), and dropout rate (0.0 to 0.2) were optimized. For the standard BINN, the learning rate (2 *×* 10*^−^*^3^ to 6 *×* 10*^−^*^3^; log-uniform), dropout rate (0.0 to 0.07), and Huber loss delta (0.025 to 1.0; log-uniform) were optimized.

### Ablation analysis

We evaluated the contributions of three architectural components by varying the ensemble size (*K* = 1 or *K* = 8), layer configuration (gated or sparse), and supervision strategy (final-only or multistage). Testing all combinations of these factors produced eight model configurations. All configurations were trained on the complete development collection using the same fixed hyperparameters, optimization settings, training duration, and early-stopping procedure. Each configuration was trained with five random seeds (1, 2, 3, 4, and 5) and evaluated on the eight held-out GEO datasets, comprising 4,431 samples (Supplementary Table S1). Performance was assessed using MSE, MAE, MedAE, and Pearson *R*, and values are reported as the mean *±* standard deviation across seeds.

### Age-associated pathway gating

Sample-specific gating values were extracted from the L2 pathway layer for the samples in the held-out testing datasets. For each sample and pathway representation, gating values were averaged across the eight ensemble members to obtain one value per sample. Each of the 504 L2 pathway representations was analyzed separately using ordinary least squares regression. Gating value was modeled as a function of chronological age, with tissue and study included as covariates. Age was standardized before model fitting, so the age coefficient represented the expected change in gating value per standard-deviation increase in age. Two-sided *P* values for the age coefficient were adjusted across the 504 representations using the Benjamini–Hochberg procedure.

To express each association on the original gating scale, the fitted model was used to estimate the change in gating value between ages 20 and 90:

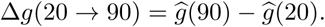

### Age-by-tissue interactions in pathway gating

We next tested whether associations between age and gating differed across tissues. For each of the 504 L2 pathway representations, gating value was modeled as a function of chronological age, tissue, study, and age-by-tissue interaction terms. The overall interaction was tested using a partial Wald *F* -test of whether all age-by-tissue interaction coefficients were jointly equal to zero. Blood tissue was the reference category, but significance was determined from the overall interaction test rather than from individual interaction coefficients. Interaction *P* values were adjusted across the 504 representations using the Benjamini-Hochberg procedure. Tissue-specific age slopes were then estimated from each fitted model. For each gene or pathway node, the extent of tissue variation was summarized as the difference between its largest and smallest tissue-specific age slopes. Among representations with an adjusted *P <* 0.05, the five with the largest slope differences were selected for presentation in Fig. 2b. No pathways were manually excluded, and no redundancy filtering was applied.

### TCGA residual epigenetic age-acceleration analysis

Within each TCGA cohort, residual epigenetic age acceleration was defined as the residual from a linear regression of predicted age on chronological age. The regression was fitted jointly across all normal and tumor samples within the cohort. Positive residuals indicated an older predicted age than expected from the cohort-specific relationship between predicted and chronological age, whereas negative residuals indicated a younger predicted age than expected.

Residual epigenetic age acceleration was compared between normal and tumor samples within each cohort using a two-sided Mann-Whitney *U* test. The comparisons were unpaired because all available normal and tumor samples were included rather than only patient-matched samples. The resulting *P* values were adjusted across the 11 cohort-level comparisons using the Benjamini-Hochberg procedure. An adjusted *P <* 0.05 was considered statistically significant. Unless otherwise stated, all statistical tests were two-sided.

### Epigenetic Clock Quantification

Published epigenetic clocks were evaluated using pyaging version 0.1.22. The benchmark included ZhangBLUP, ZhangEN, PCHannum, Hannum, YingAdaptAge, YingDamAge, and YingCausAge. Predictions were generated using the package’s default implementation of each clock. Missing CpG values were imputed internally using the clock-specific reference values provided by pyaging. No clock-specific settings were modified.

### SHAP feature-attribution analysis

SHAP was used to estimate how gene and pathway representations contributed to EG-BINN predictions. For each hierarchy level, the representations at that level served as SHAP features, and attribution was performed for the predictions generated by the corresponding head and all down-stream heads.

For hierarchy level *ℓ*, the prediction component explained by SHAP was defined as

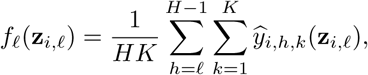

where **z***_i,ℓ_* denotes the representation of sample *i* at level *ℓ*, *H* = 4 is the total number of prediction heads, and *K* = 8 is the number of ensemble members. Each analysis therefore explained the prediction component generated at or downstream of the selected hierarchy level.

Attributions were estimated using shap.GradientExplainer. SHAP values were first calculated for the member-specific representations and then summed across the eight ensemble members to obtain one attribution per representation and sample while preserving the scale of the corresponding prediction component. For each of the six TCGA cohorts with a significant tumor–normal difference in residual epigenetic age acceleration after Benjamini–Hochberg correction, normal samples served as the SHAP background reference and tumor samples were explained. Backgrounds were defined separately for each cohort and hierarchy level using up to 100 normal samples.

The primary cross-cohort analysis focused on the 504 L2 pathway representations. Gene-, L1-, and L0-level analyses are reported in the Supplementary Information. Within each cohort, L2 representations were ranked by their mean absolute SHAP value across tumor samples, and the 20 highest-ranked representations were selected. Mean signed SHAP values were calculated separately to summarize whether each representation tended to increase or decrease predictions relative to the cohort-specific normal background. Because each cohort used a different background distribution, cross-cohort comparisons were based on within-cohort ranks and recurrence among the top 20 representations rather than on raw SHAP magnitudes.

### Use of artificial intelligence

During the preparation of this manuscript, the authors used Microsoft Copilot solely for grammatical correction and improving readability of the manuscript. In addition, we used GLM 5.2/5.3 to assist with various programming tasks for analyses, such as figure generation and code review. The authors take full responsibility for the implementation, analyses, manuscript content, and reported findings.

## Data availability

All datasets analyzed in this study are publicly available through GEO and TCGA. Details of all studies used for model training, validation, and testing are provided in Supplementary Datasets S1 and S2. Dataset information is available in our supplementary files as Supplementary Information Dataset S1 and S2. See our GitHub repository (https://github.com/Travyse/egbinn) for information on how to download the training and inference data.

## Code availability

All code used for model development, hyperparameter optimization, feature attribution analyses, and figure generation is available at our GitHub repository (https://github.com/Travyse/egbinn). Documentation and example workflows are provided to facilitate reproduction of the results presented in this study.

## Acknowledgements

This work was supported in part by the National Institutes of Health grants T32 HG000046, U01 AG068057, U19 AG074879, R01 AG071470, U01 AG066833.

## Author contributions

T.A.E. conceived the study, developed the model, designed and conducted the experiments, performed the analyses, interpreted the results, and wrote and revised the manuscript. Q.L. and L.S. reviewed and edited the manuscript. All authors read and approved the final manuscript.

## References

1. World Health Organization. Ageing and health. Accessed: 2026-09-21. 2025. url: https://www.who.int/news-room/fact-sheets/detail/ageing-and-health.

2. Horvath S. DNA methylation age of human tissues and cell types. Genome Biology 2013;14.

3. Bernabeu E, McCartney DL, Gadd DA, et al. Refining epigenetic prediction of chronological and biological age. Genome Medicine 2023;15.

4. Levine ME, Lu AT, Quach A, et al. An epigenetic biomarker of aging for lifespan and healthspan. Aging 2018;10:573–91.

5. Lu AT, Quach A, Wilson JG, et al. DNA methylation GrimAge strongly predicts lifespan and healthspan. Aging 2019;11:303–27.

6. Lu AT, Binder AM, Zhang J, et al. DNA methylation GrimAge version 2. Aging 2022.

7. Johnson ND, Wiener HW, Smith AK, et al. Non-linear patterns in age-related DNA methylation may reflect CD4^+^ T cell differentiation. Epigenetics 2017;12:492–503.

8. Olecka M, Bommel A van, Best L, et al. Nonlinear DNA methylation trajectories in aging male mice. Nature Communications 2024;15.

9. Galkin F, Mamoshina P, Kochetov K, Sidorenko D, and Zhavoronkov A. DeepMAge: A Methylation Aging Clock Developed with Deep Learning. Aging and disease 2021;12:1252.

10. Levy JJ, Titus AJ, Petersen CL, Chen Y, Salas LA, and Christensen BC. MethylNet: an automated and modular deep learning approach for DNA methylation analysis. BMC Bioinformatics 2020;21.

11. Lima Camillo LP de, Lapierre LR, and Singh R. A pan-tissue DNA-methylation epigenetic clock based on deep learning. npj Aging 2022;8.

12. Lin A, Giosan I, Aparicio A, et al. DeepStrataAge: an interpretable deep-learning clock that reveals stage- and sex-divergent DNA methylation aging dynamics. npj Aging 2026;12.

13. Prosz A, Pipek O, Borcsok J, et al. Biologically informed deep learning for explainable epigenetic clocks. Scientific Reports 2024;14.

14. Greene CS, Krishnan A, Wong AK, et al. Understanding multicellular function and disease with human tissue-specific networks. Nature Genetics 2015;47:569–76.

15. Sonawane AR, Platig J, Fagny M, et al. Understanding Tissue-Specific Gene Regulation. Cell Reports 2017;21:1077–88.

16. Aguet F, Anand S, Ardlie KG, et al. The GTEx Consortium atlas of genetic regulatory effects across human tissues. Science 2020;369:1318–30.

17. Jayakumar SM, Menick J, Czarnecki WM, et al. Multiplicative Interactions and Where to Find Them. In: International Conference on Learning Representations. 2020. url: https://api.semanticscholar.org/CorpusID:213009174.

18. Dauphin YN, Fan A, Auli M, and Grangier D. Language Modeling with Gated Convolutional Networks. 2016.

19. Shazeer N. GLU Variants Improve Transformer. 2020.

20. Gorishniy Y, Kotelnikov A, and Babenko A. TabM: Advancing Tabular Deep Learning with Parameter-Efficient Ensembling. 2024.

21. Conard AM, DenAdel A, and Crawford L. A spectrum of explainable and interpretable machine learning approaches for genomic studies. WIREs Computational Statistics 2023;15.

22. Marcinkevics R and Vogt JE. Interpretable and explainable machine learning: A methodscentric overview with concrete examples. WIREs Data Mining and Knowledge Discovery 2023;13.

23. Hartman E, Scott AM, Karlsson C, et al. Interpreting biologically informed neural networks for enhanced proteomic biomarker discovery and pathway analysis. Nature Communications 2023;14.

24. Elmarakeby HA, Hwang J, Arafeh R, et al. Biologically informed deep neural network for prostate cancer discovery. Nature 2021;598:348–52.

25. Hanahan D. Hallmarks of Cancer: New Dimensions. Cancer Discovery 2022;12:31–46.

26. Hanahan D and Monje M. Cancer hallmarks intersect with neuroscience in the tumor microenvironment. Cancer Cell 2023;41:573–80.

27. Hanahan D. Hallmarks of cancer—Then and now, and beyond. Cell 2026;189:2254–77.

28. Ragueneau E, Gong C, Sinquin P, et al. The Reactome Knowledgebase 2026. Nucleic Acids Research 2025;54:D673–D681.

29. Clough E and Barrett T. The Gene Expression Omnibus Database. In: Methods in Molecular Biology. Springer New York, 2016:93–110. doi: 10.1007/978-1-4939-3578-9_5.

30. Weinstein JN, Collisson EA, Mills GB, et al. The Cancer Genome Atlas Pan-Cancer analysis project. Nature Genetics 2013;45:1113–20.

