## Supplementary Information for "EG-BINN: Parameter-Efficient Ensembling of Gated Biologically Informed Neural Networks for Epigenetic Age Estimation"

#### This PDF file includes:

Figs. S1 to S10
Tables S1 to S3
Legends for Dataset S1 to S2

#### Other supporting materials for this manuscript include the following:

Datasets S1 to S2

**Table S1. Ablation analysis of EG-BINN architectural components**

| K | Layer Type | Regression | Final Configuration | Pearson R ↑ | MAE ↓ | MedAE ↓ | MSE ↓ |
|---|---|---|---|---|---|---|---|
| 8 | Gated | Multistage | ✓ | **0.86 ± 0.01** | **4.03 ± 0.10** | **2.51 ± 0.10** | **38.61 ± 1.96** |
| 1 | Gated | Multistage | × | 0.85 ± 0.01 | 4.10 ± 0.08 | 2.56 ± 0.05 | 40.30 ± 2.23 |
| 8 | Sparse | Multistage | × | 0.83 ± 0.01 | 4.26 ± 0.12 | 2.60 ± 0.06 | 44.33 ± 2.54 |
| 1 | Sparse | Multistage | × | 0.83 ± 0.01 | 4.35 ± 0.18 | 2.67 ± 0.12 | 45.79 ± 3.30 |
| 8 | Gated | Final | × | 0.83 ± 0.02 | 4.45 ± 0.19 | 2.85 ± 0.11 | 45.11 ± 4.66 |
| 1 | Gated | Final | × | 0.79 ± 0.02 | 5.10 ± 0.29 | 3.28 ± 0.27 | 57.00 ± 4.40 |
| 8 | Sparse | Final | × | 0.77 ± 0.01 | 5.22 ± 0.17 | 3.29 ± 0.19 | 62.15 ± 1.59 |
| 1 | Sparse | Final | × | 0.76 ± 0.02 | 5.39 ± 0.34 | 3.48 ± 0.27 | 64.18 ± 5.23 |

**Table S2. Parameter counts and structural connectivity across model architectures**

| Metric | BINN | EG-BINN | MLP |
|---|---|---|---|
| Total params | 278,629 | 2,692,468 | 24,246,529 |
| Input dim (gene-mapped) | 189,291 | 189,291 | 189,291 |
| CpG→gene edges | 209,839 | 209,839 (×2: value + gate) | — |
| Connected genes | 20,031 | 20,031 | — |
| First-layer params | 229,870 | 2,294,564 | 24,229,632 |
| First-layer % of total | 82.5% | 85.2% | 99.93% |
| Dense CpG→gene equivalent | 3,791,708,052 | 3,791,708,052 | — |

**Table S3. Parameter efficiency of BatchEnsemble sharing in EG-BINN**

| Configuration | Members | Shared base | Per-member | Total |
|---|---|---|---|---|
| Non-ensembled gated BINN | 1 | – | 536,588 | 536,588 |
| $K = 8$ EG-BINN (shared base) | 8 | 515,860 | 272,076 | 2,692,468 |
| 8 × non-ensembled (independent) | 8 | – | 536,588 | 4,292,704 |

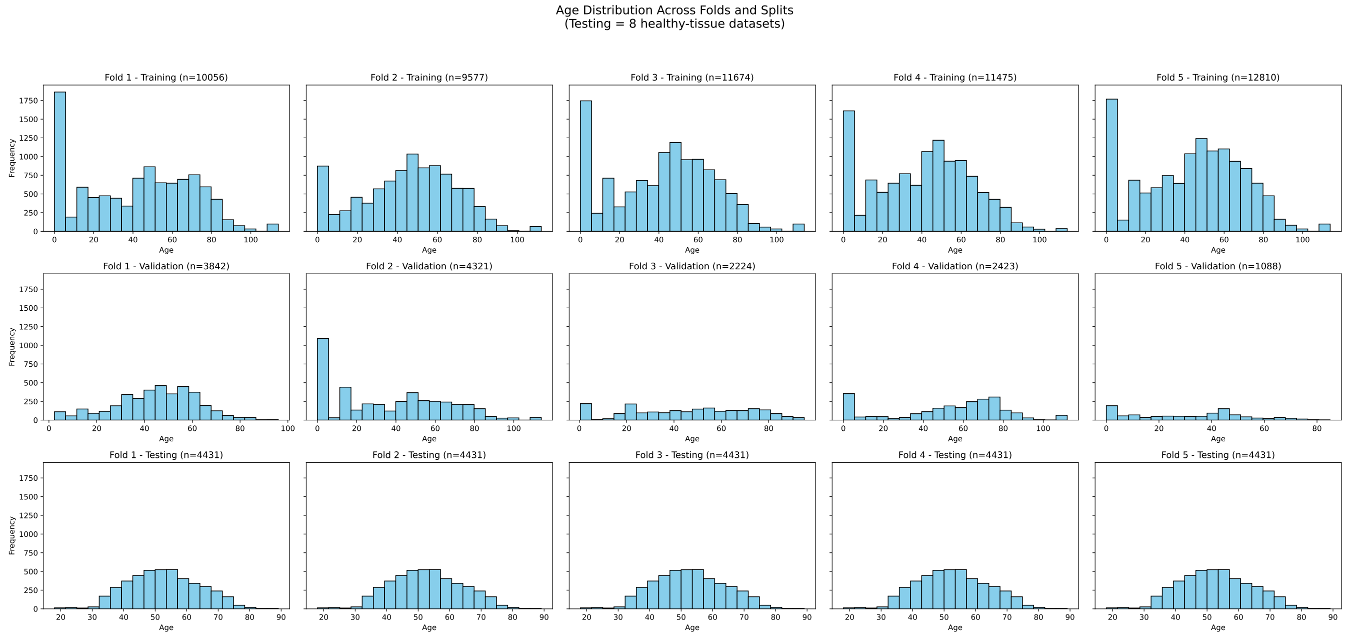


**Fig. S1.** Age distributions across cross-validation folds and data splits. Histograms show chronological-age distributions for training (top row), validation (middle row), and held-out evaluation samples (bottom row) for each of the five dataset-level cross-validation folds. Samples from 93 GEO datasets measured using the Illumina HumanMethylation450K or MethylationEPIC platforms were assigned to folds at the dataset level, ensuring that all samples from a study remained within the same partition. A fixed set of eight GEO datasets was excluded before fold construction and retained for held-out evaluation in every fold. Each panel is annotated with the corresponding sample count, and rows share a common $y$-axis.

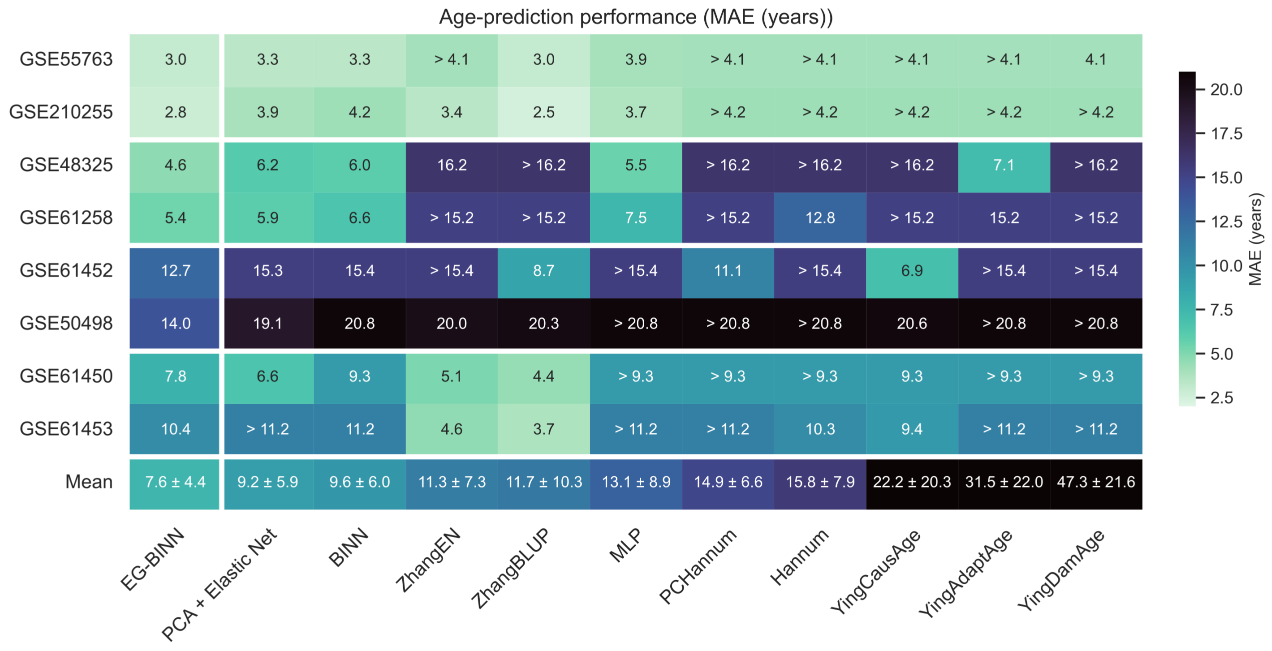


**Fig. S2.** Mean absolute error of each aging model in the eight held-out testing datasets. Columns are ordered by increasing mean cohort-level MAE. The summary row reports the unweighted mean ± standard deviation across cohorts.

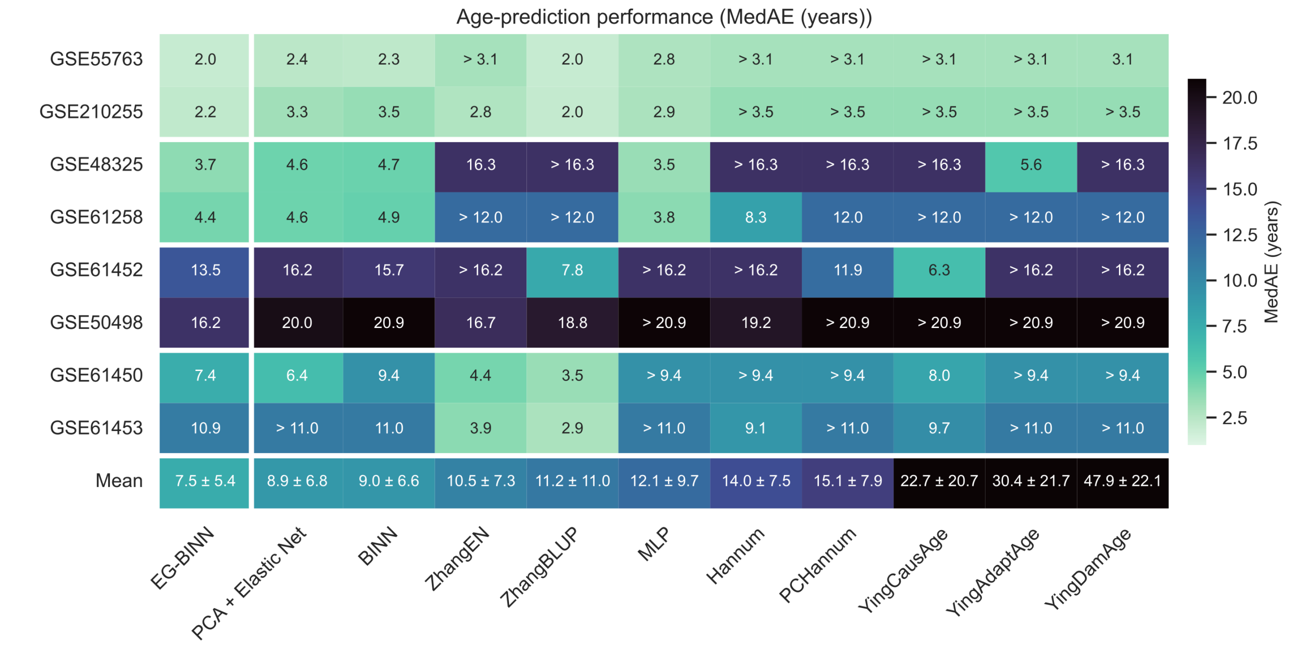


**Fig. S3.** Median absolute error of each aging model in the eight held-out testing datasets. Columns are ordered by increasing mean cohort-level MedAE. The summary row reports the unweighted mean ± standard deviation across cohorts.

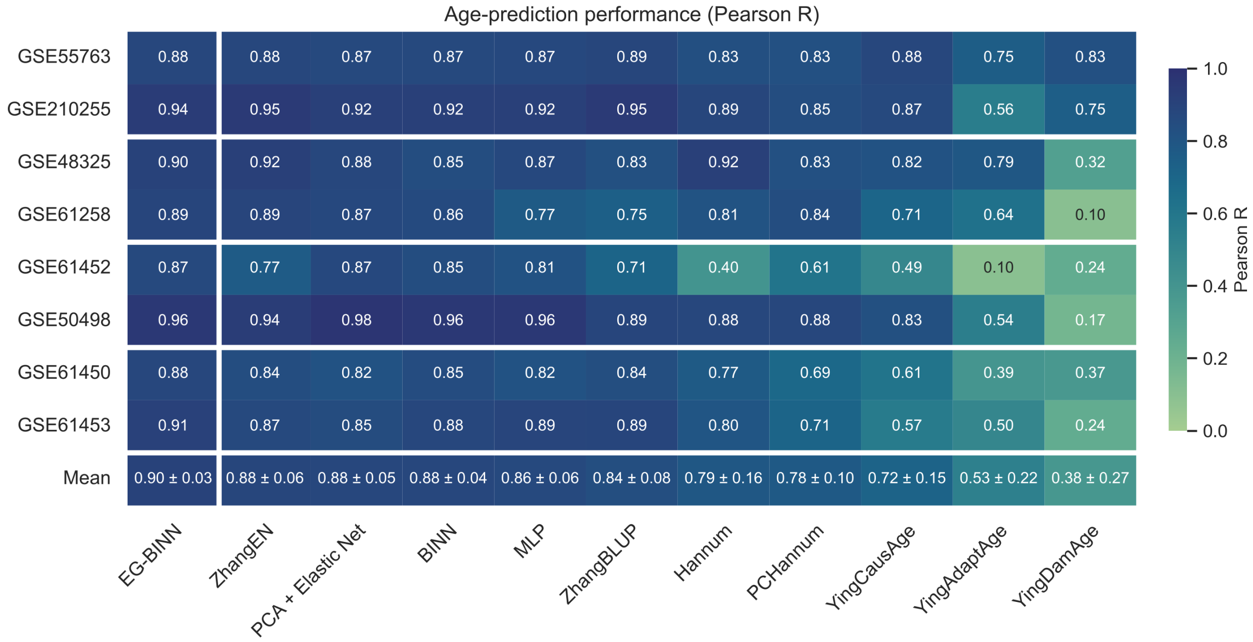


**Fig. S4.** Pearson R of each aging model in the eight held-out testing datasets. Columns are ordered by decreasing mean cohort-level Pearson R. The summary row reports the unweighted mean ± standard deviation across cohorts.

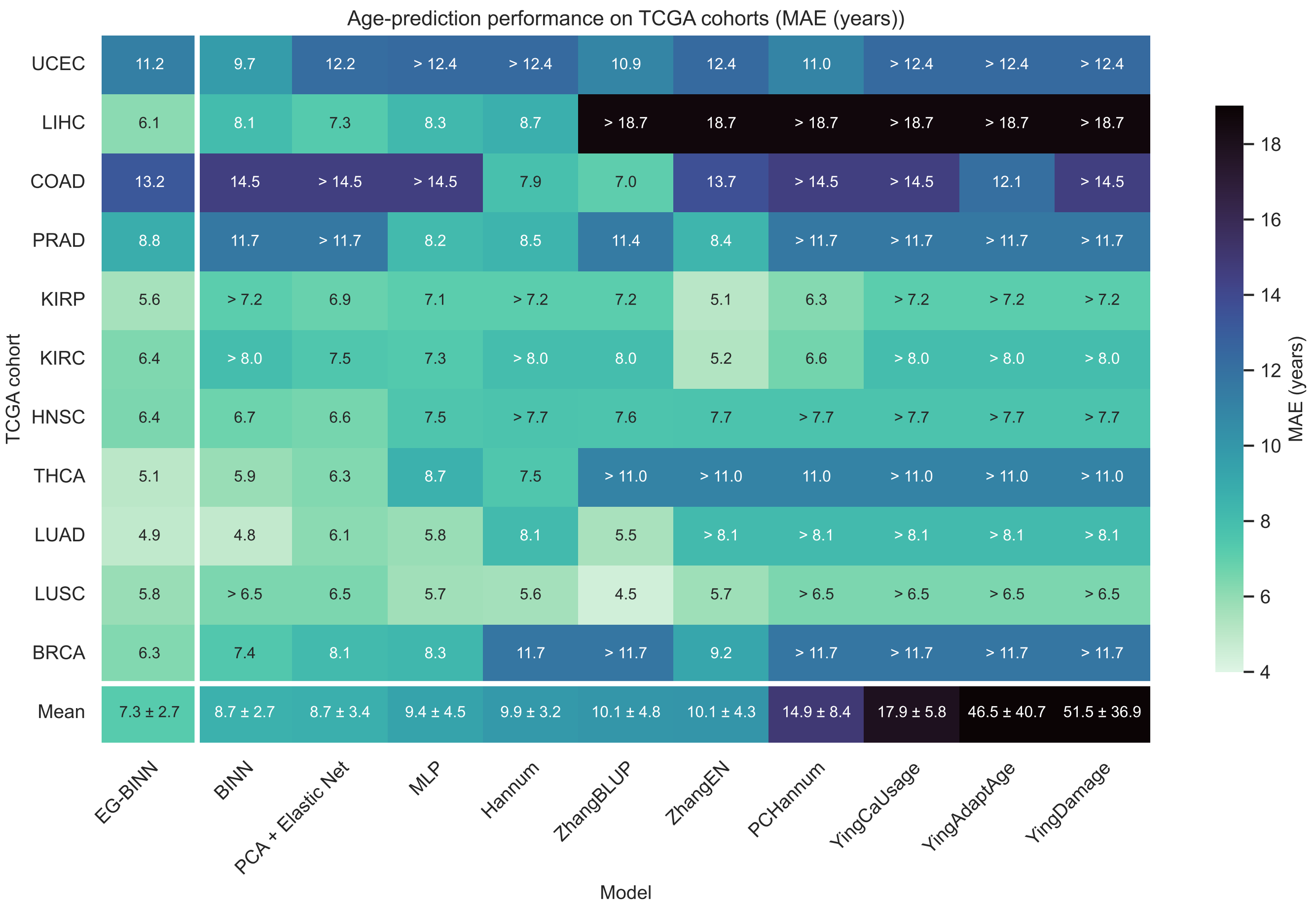


**Fig. S5.** Mean absolute error of each aging model in normal tissue samples from all 11 TCGA cohorts. Columns are ordered by increasing mean cohort-level MAE. The summary row reports the unweighted mean ± standard deviation across cohorts.

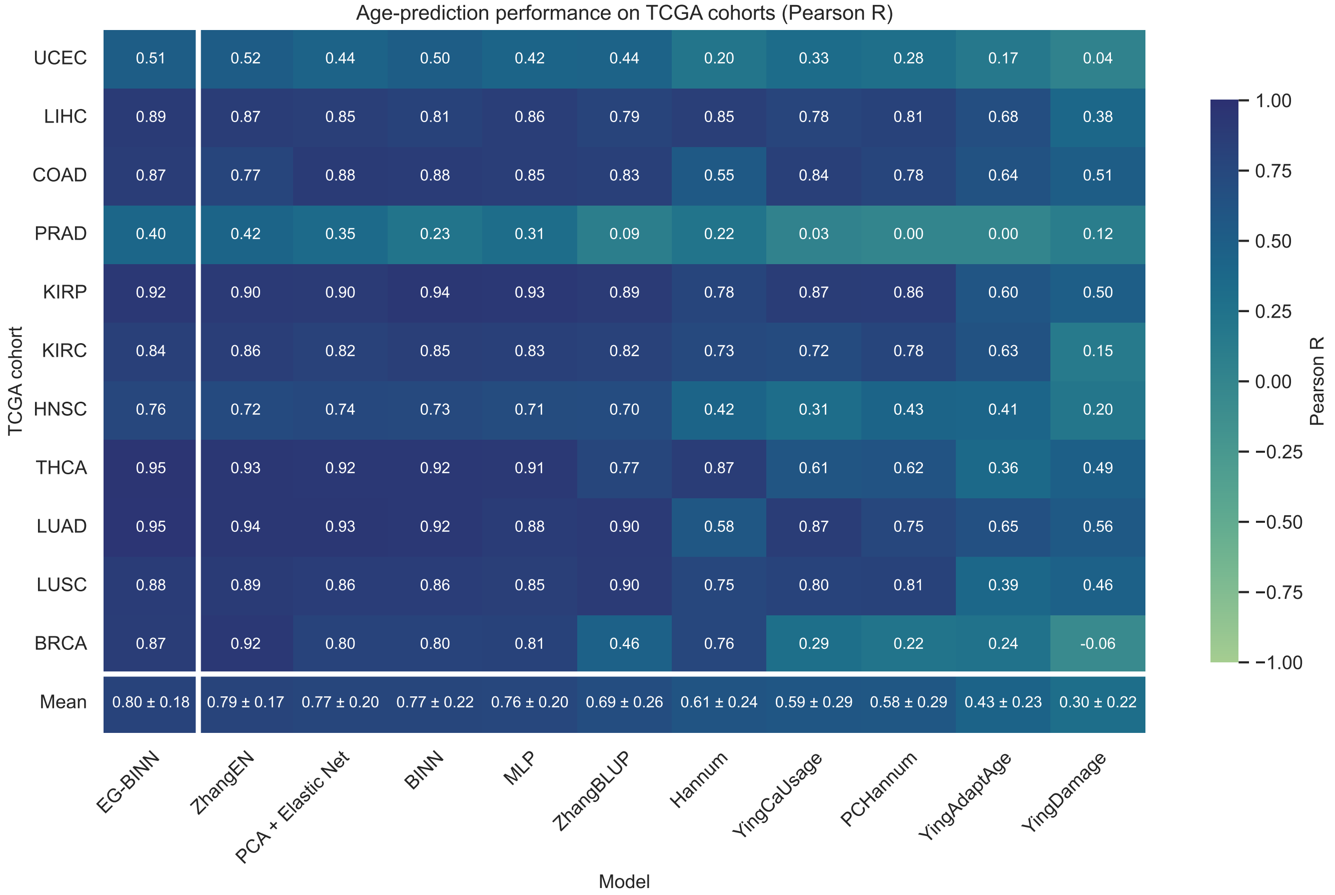


**Fig. S6.** Pearson R of each aging model in normal tissue samples from all 11 TCGA cohorts. Columns are ordered by decreasing mean cohort-level Pearson R. The summary row reports the unweighted mean ± standard deviation across cohorts.

 

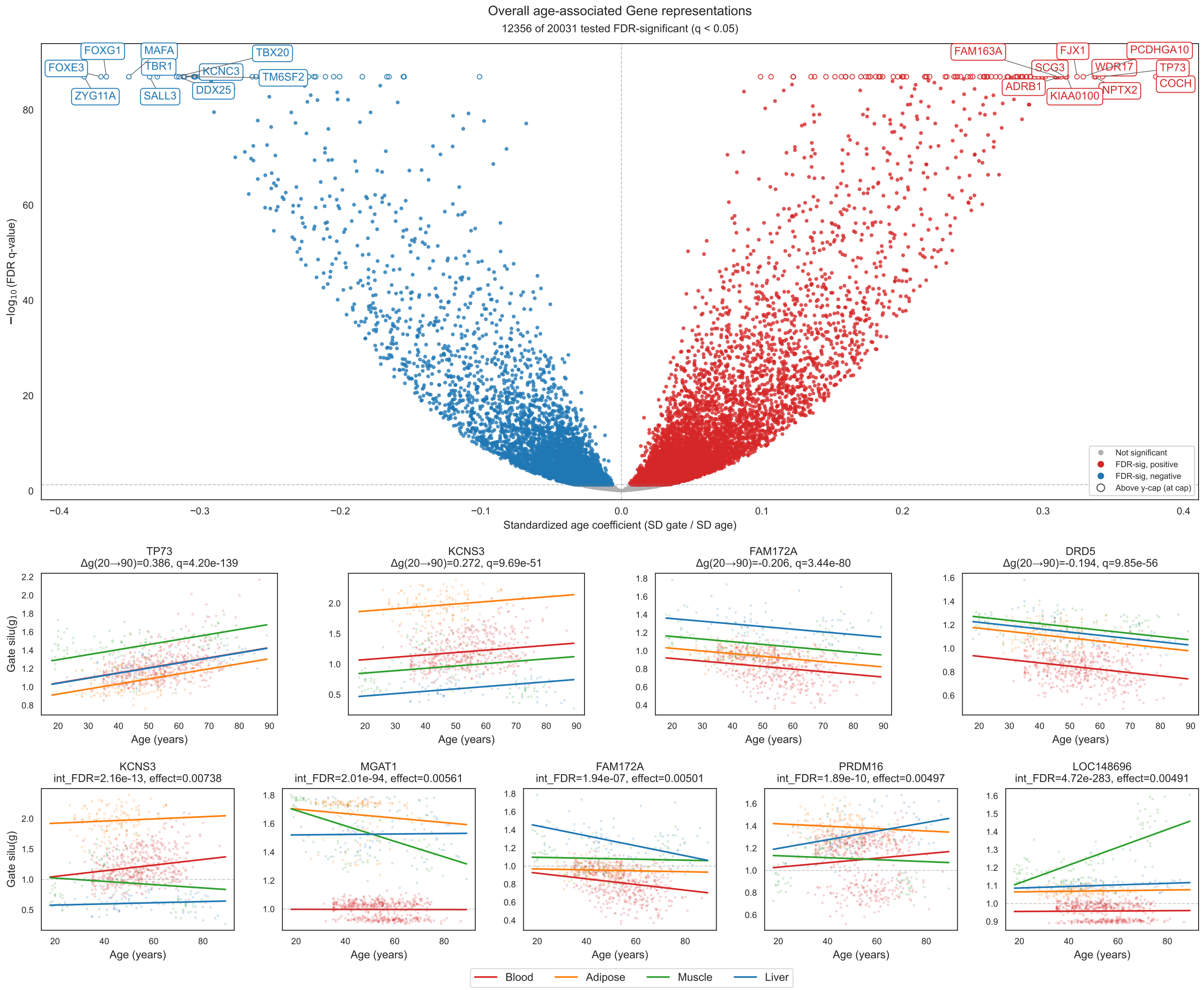


**Fig. S7. Age- and tissue-dependent modulation of gene representations in EG-BINN. a**, Standardized associations between chronological age and gating values for 20,031 gene representations, estimated across blood, adipose tissue, skeletal muscle, and liver with adjustment for tissue and study. Filled points indicate associations significant after Benjamini–Hochberg correction. Trajectory plots show adjusted gating values for the significant representations with the largest positive and negative estimated changes from ages 20 to 90. **b**, Adjusted tissue-specific trajectories for five gene representations with significant age-by-tissue interactions and the largest differences among tissue-specific age slopes. Lines show fitted values, and shaded bands indicate 95% confidence intervals. The horizontal dashed line at a gate value of 1 indicates neutral modulation; values above 1 indicate amplification, whereas values between 0 and 1 indicate attenuation of the corresponding representation.

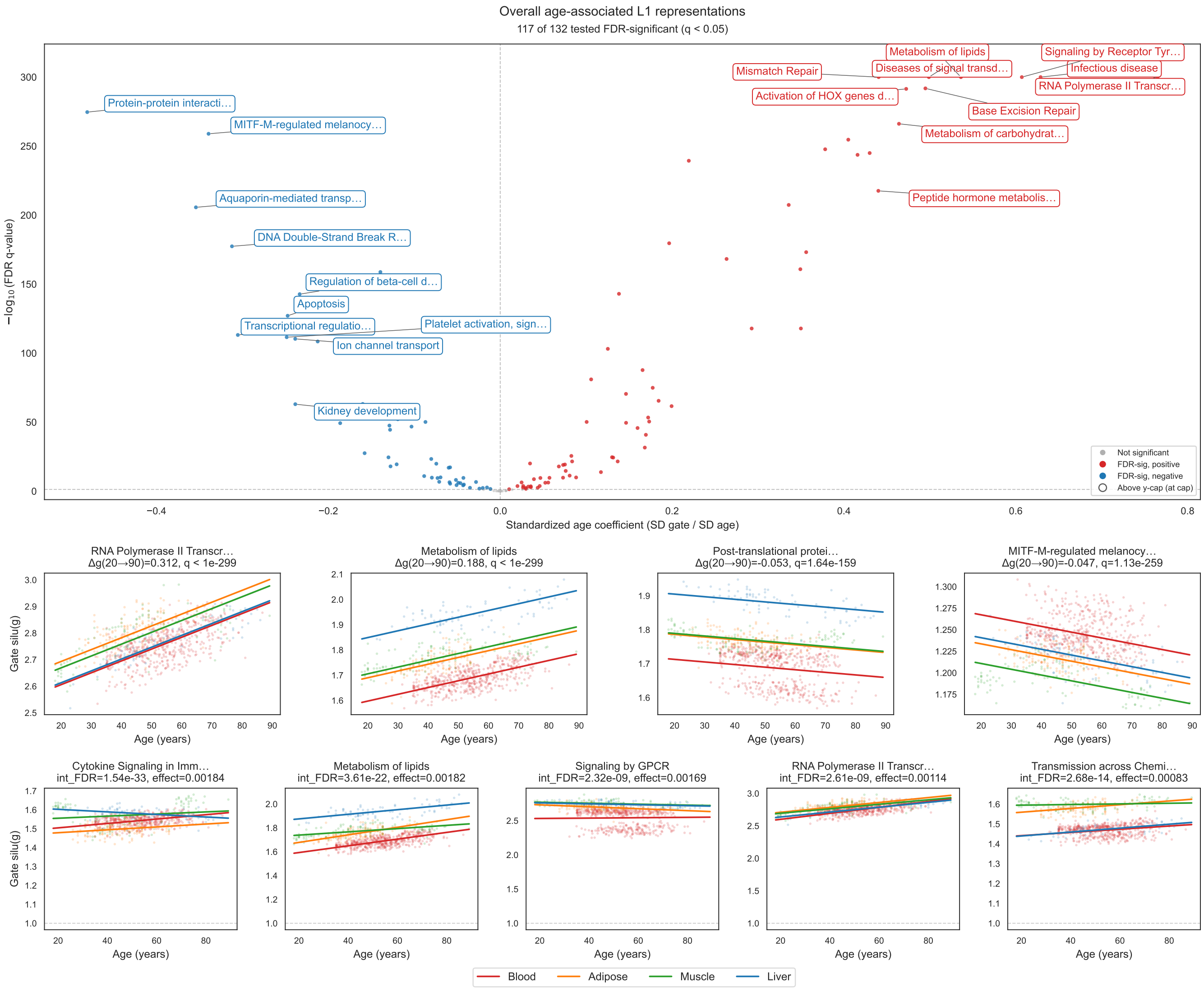


**Fig. S8. Age- and tissue-dependent modulation of L1 pathway representations in EG-BINN. a**, Standardized associations between chronological age and gating values for 132 L1 pathway representations, estimated across blood, adipose tissue, skeletal muscle, and liver with adjustment for tissue and study. Filled points indicate associations significant after Benjamini–Hochberg correction. Trajectory plots show adjusted gating values for the significant representations with the largest positive and negative estimated changes from ages 20 to 90. **b**, Adjusted tissue-specific trajectories for five L1 pathway representations with significant age-by-tissue interactions and the largest differences among tissue-specific age slopes. Lines show fitted values, and shaded bands indicate 95% confidence intervals. The horizontal dashed line at a gate value of 1 indicates neutral modulation; values above 1 indicate amplification, whereas values between 0 and 1 indicate attenuation of the corresponding representation.

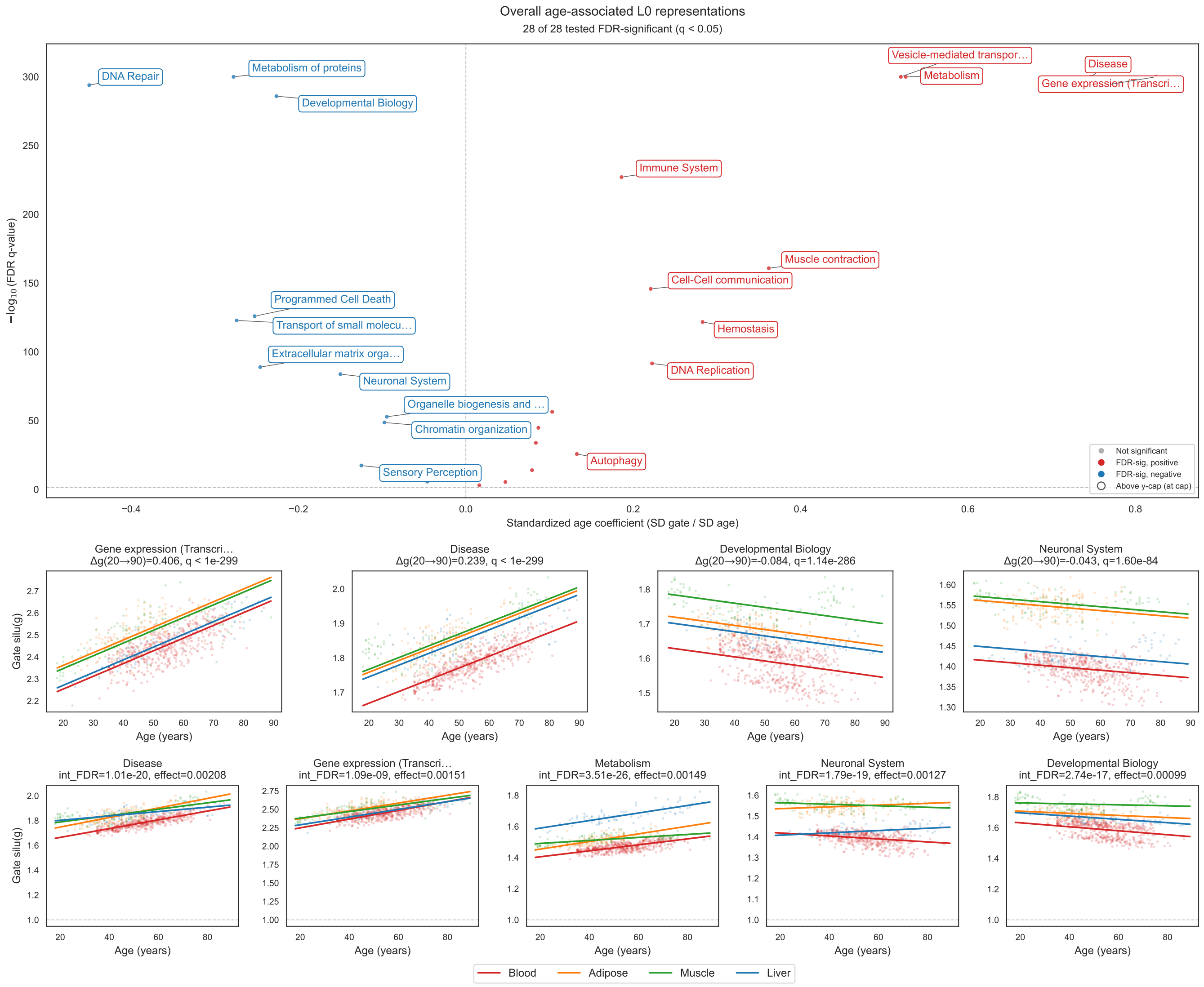


**Fig. S9. Age- and tissue-dependent modulation of L0 pathway representations in EG-BINN. a**, Standardized associations between chronological age and gating values for 28 L0 pathway representations, estimated across blood, adipose tissue, skeletal muscle, and liver with adjustment for tissue and study. Filled points indicate associations significant after Benjamini–Hochberg correction. Trajectory plots show adjusted gating values for the significant representations with the largest positive and negative estimated changes from ages 20 to 90. **b**, Adjusted tissue-specific trajectories for five L0 pathway representations with significant age-by-tissue interactions and the largest differences among tissue-specific age slopes. Lines show fitted values, and shaded bands indicate 95% confidence intervals. The horizontal dashed line at a gate value of 1 indicates neutral modulation; values above 1 indicate amplification, whereas values between 0 and 1 indicate attenuation of the corresponding representation.

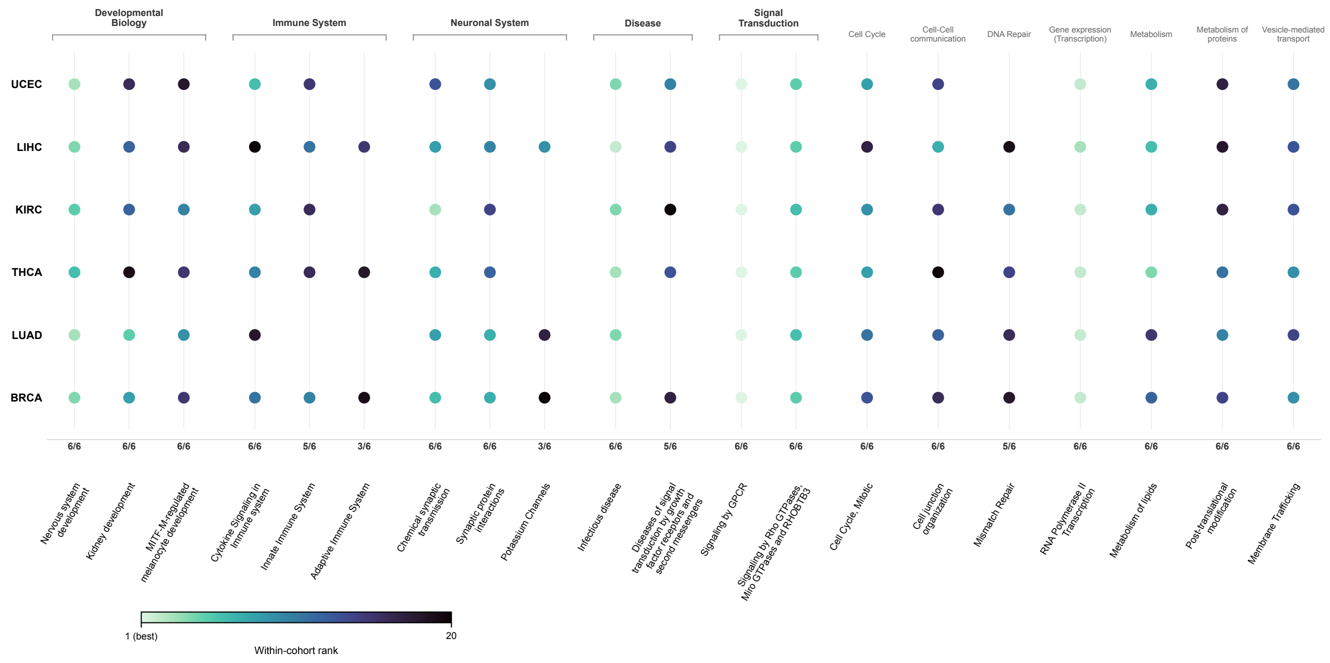


**Fig. S10.** L1-level SHAP attribution for tumor samples using cohort-specific normal samples as the background reference distribution. Rows represent the six analyzed TCGA cohorts, and columns represent L1 pathways retained among the 20 highest-ranked representations in at least three cohorts. Within each cohort, pathways were ranked according to mean absolute SHAP value across tumor samples. Dot color denotes within-cohort rank, with lighter colors indicating higher-ranked pathways, and dot size is uniform. A missing dot indicates that the pathway was not retained among the corresponding cohort's top 20 L1 pathways. Values beneath pathway columns indicate the number of cohorts in which each pathway was retained. Pathway attributions quantify contributions to downstream multistage prediction components and do not directly measure pathway activation or causal involvement in residual epigenetic age acceleration.

### SI Dataset S1 (geo_data_supplementary_file.csv)

Dataset-level characteristics of the 93 development datasets and eight held-out evaluation datasets. The file reports GEO accession, methylation platform, primary tissue, sample counts, chronological-age summary statistics (minimum, maximum, mean, and standard deviation), and development or held-out assignment. All samples from a given GEO dataset were assigned to the same cross-validation fold to prevent study-level leakage. The eight held-out datasets were excluded before construction of the cross-validation folds and were used for the held-out benchmark evaluation.

### SI Dataset S2 (tcga_supplementary_file.csv)

Cohort- and tissue-class-level characteristics of the 11 TCGA cohorts evaluated in this study, with one row per cohort and tissue class (tumor or normal). The file reports TCGA cohort identifier, cancer name, tissue class, methylation platform, sample counts, and chronological-age summary statistics (minimum, maximum, mean, standard deviation, median, and interquartile range). Samples with missing age values were excluded prior to aggregation.
